# Regeneration of spinal motor axons: Negative regulation by Celsr2 implicating Cdc42/Rac1 and JNK/c-Jun signaling

**DOI:** 10.1101/2021.01.20.427520

**Authors:** Quan Wen, Huandi Weng, Tao Liu, Lingtai Yu, Tainyun Zhao, Jingwen Qin, Tissir Fadel, Yibo Qu, Libing Zhou

## Abstract

During development, cadherins Celsr2 and Celsr3 control axon navigation. Unlike Celsr3, Celsr2 remains expressed in the adult, suggesting unexplored roles in maintenance and repair. Here we show that *Celsr2* knockdown promotes motor axon regeneration in mouse and human spinal cord explants and cultured motor neurons. *Celsr2* downregulation is accompanied by increased levels of GTP-bound Rac1 and Cdc42, and of JNK and c-Jun proteins. Using a branchial plexus injury model, we show that forelimb functional recovery is improved in *Celsr2* mutant versus control mice. Compared to controls, in mutant mice, reinnervated biceps muscles are less atrophic, contain more newly formed neuromuscular junctions, and generate larger electromyographic potentials, while motor neuron survival and axon regeneration are improved. GTP-bound Rac1 and Cdc42, JNK and c-Jun are upregulated in injured mutant versus control spinal cord. In conclusion, Celsr2 negatively regulates motor axon regeneration via Cdc42/Rac1/JNK/c-Jun signaling and is a target for neural repair.

## Introduction

Spinal motor neurons are unique in that their somas and dendrites reside in the central nervous system (CNS) whereas their axons extend in the peripheral nervous system (PNS), giving them qualities of both CNS and PNS neurons. For instance, following injury, spinal motor axons regenerate better than those of premotor CNS neurons (Liu, Tedeschi, Park, & He, 2011; Mahar & Cavalli, 2018). Brachial plexus injury (BPI) often leads to root avulsion and lesions of motor neurons and axons, with dramatic consequences, and represents an important health issue. Root lesions interrupt corresponding segmental motor axons and sensory fibers, leading to axon degeneration and, eventually, death of motor neurons (Ruven, Chan, & Wu, 2014). Contrary to distal motor axon injuries, axons regenerate poorly after root avulsion. Reimplantation of avulsed roots in the spinal cord provides a scaffold that may facilitate survival of injured motor neurons and regeneration of their axons (Cullheim, Carlstedt, Linda, Risling, & Ulfhake, 1989). These observations resulted in development of a surgical method to restore function after root injury (Carlstedt, Grane, Hallin, & Noren, 1995). Potential therapies to promote axon regrowth and functional recovery have been considered (Duraikannu, Krishnan, Chandrasekhar, & Zochodne, 2019; Havton & Carlstedt, 2009). Yet, the limits of current treatments outline the need for further basic research into mechanisms that enhance motor axon regeneration (Carlstedt, 2010).

Together with its two paralogs Celsr1 and Celsr3, the atypical cadherin Celsr2 is an orthologue of the *Drosophila* planar cell polarity (PCP) protein Flamingo. Celsr1-3 play critical roles during brain development (Goffinet & Tissir, 2017; Tissir & Goffinet, 2013). Genetic studies in mice showed that Celsr2 and Celsr3 regulate axonal pathfinding, motoneuron migration and ependymal ciliogenesis (Chai et al., 2014; Qu et al., 2010; Qu et al., 2014; Tissir et al., 2010). Whereas mutant phenotypes suggest that Celsr2 and Celsr3 act synergistically, they were also reported to play opposite roles in steering neurite outgrowth *in vitro* (Shima et al., 2007).

Motor axon phenotypes observed in *Celsr3* and *Celsr2* double mutant mice (Chai et al., 2014) are also observed in *Fzd3* mutant mice (Hua, Smallwood, & Nathans, 2013) and upon motor neuron-specific inactivation of *Rac1* (Hua, Emiliani, & Nathans, 2015), hinting at a mechanistic link between Celsr2 & 3, Fzd3 and GTPase-dependent regulation of growth cone extension. Contrary to *Celsr3*, the expression of which is sharply downregulated after birth, *Celsr2* mRNA expression remains high in the adult CNS (Tissir & Goffinet, 2006). This prompted us to investigate whether Celsr2 has any effect on the regeneration of injured motor axons.

To address this, we compared motor axon outgrowth in control and *Celsr2*-deficient spinal cord explants and cultured motor neurons from mouse and human embryos.

Human *CELSR2* was downregulated using a lentivirus encoding *Celsr2* shRNA, whereas mouse *Celsr2* was inactivated genetically. To investigate the role of Celsr2 during adult motor axon regeneration, we performed spinal root avulsion and motor root reimplantation in control and *Celsr2* motor neuron-specific knockout mice, and evaluated axon regeneration and functional recovery after surgery.

## Results

### Celsr2 is strongly expressed in spinal motor neurons

To study Celsr2 expression, we used *Celsr2*^*LacZ/+*^ mice, in which an internal ribosomal entry site (IRES) and the *LacZ* gene are inserted into *Celsr2* exon 23 (Tissir et al., 2010). Celsr2 expressing cells were visualized by anti-*β*-gal staining. *β*-gal signal was found in several cell types (Figure 1, A and F). In E12.5 spinal sections, all Isl1-positive cells in the ventral horn co-expressed *β*-gal (Figure 1, A-E). In adult spinal sections, motor neurons were positive using anti-choline acetyl transferase (ChAT) immunostaining and all of them were also positive for *β*-gal (Figure 1, F-J). Thus, Celsr2 is strongly expressed in embryonic and adult spinal motor neurons.

**Figure 1.**
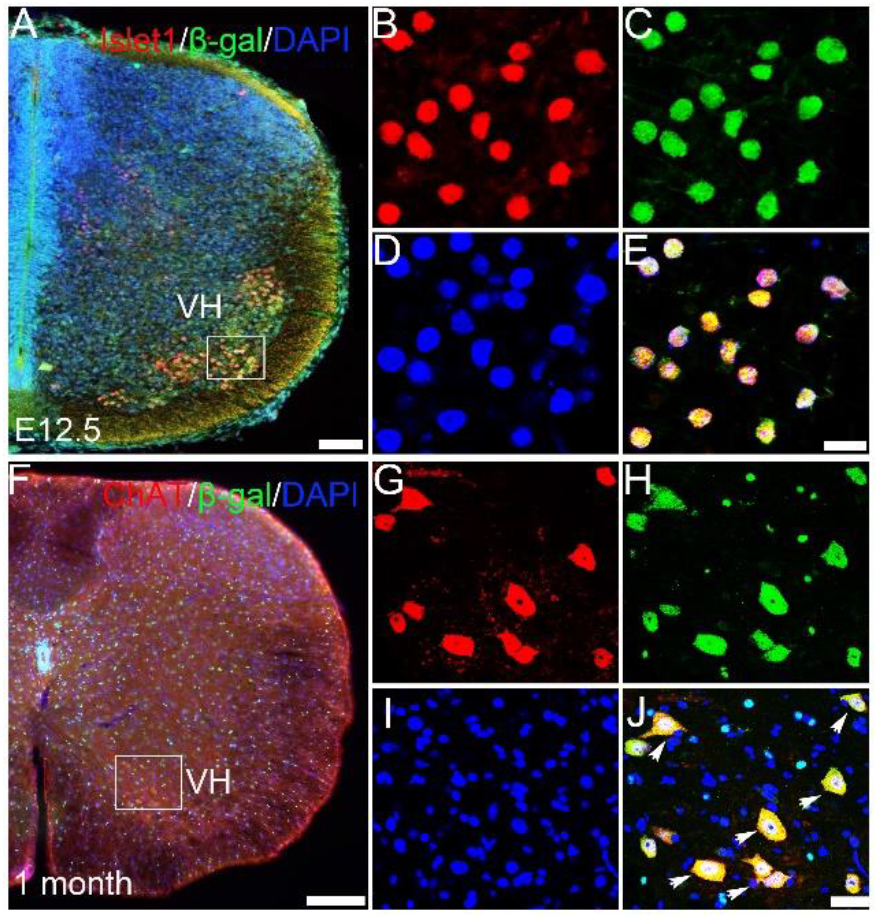
Celsr2 expression is enriched in embryonic and adult spinal motor neurons. Using *Celsr2*^*LacZ*^ transgenic mice, Celsr2 expression is detected by anti-*β*-gal immunostaining. (A-E) E12.5 spinal sections were immunostained for Islet1 (red), *β*-gal (green) and counterstained for DAPI (blue). The merged image showed that all Islet1-positive cells in the ventral horn (VH) co-labeled by *β*-gal immunoreactivity (E). (F-J) In adult spinal sections, all ChAT-positive (red) spinal motor neurons co-expressed *β*-gal (green). Nuclei were counterstained by DAPI (blue). B-E and G-J correspond to boxed areas in A and F respectively. Scale bars: 50 µm in A, 10 µm in B-E, 200 µm in F, 50 µm in G-J.

### Inactivation of *Celsr2* promotes axon growth and growth cone expansion in mouse spinal motor neurons

To assess the role of Celsr2 in motor axon regeneration, we used spinal motor neuron explant cultures (Wang & Marquardt, 2012). Cervical spinal explants containing motor neurons were prepared from E13.5 *Celsr2^−/−^* and control (*Celsr2*^*+/+*^) embryos and cultured for 6 days *in vitro* (DIV, Figure 2A). Cells and outgrowing axons expressed ChAT in both groups (Figure 2, B and E), indicating that they corresponded to spinal motor neurons from the ventral column. Anti-Tuj1 signal colocalized extensively with ChAT-immunoreactivity (Figure 2, C, D, F and G). In Tuj1-immunostained preparations, the maximal area covered by growing axons, and axonal length (indicated in Figure 2A) were significantly increased in *Celsr2^−/−^* mutant compared to the control samples (Figure 2, H and I; 3 litters of embryos in each group, 5-7 embryos in each litter, 19 and 22 explants in mutant and control, respectively; *P*<0.05 and 0.01). In addition, larger axonal bundles were formed in the mutant explants (indicated in insets of Figure 2, C and F). In the region 150-200 µm away from explant borders, the number of large (>8 µm) axon bundles, estimated as described (Jaworski & Tessier-Lavigne, 2012), was significantly higher in mutant explants (Figure 2J; *P*<0.001). These results suggest that inactivation of *Celsr2* stimulates spinal motor axon growth and fasciculation.

**Figure 2.**
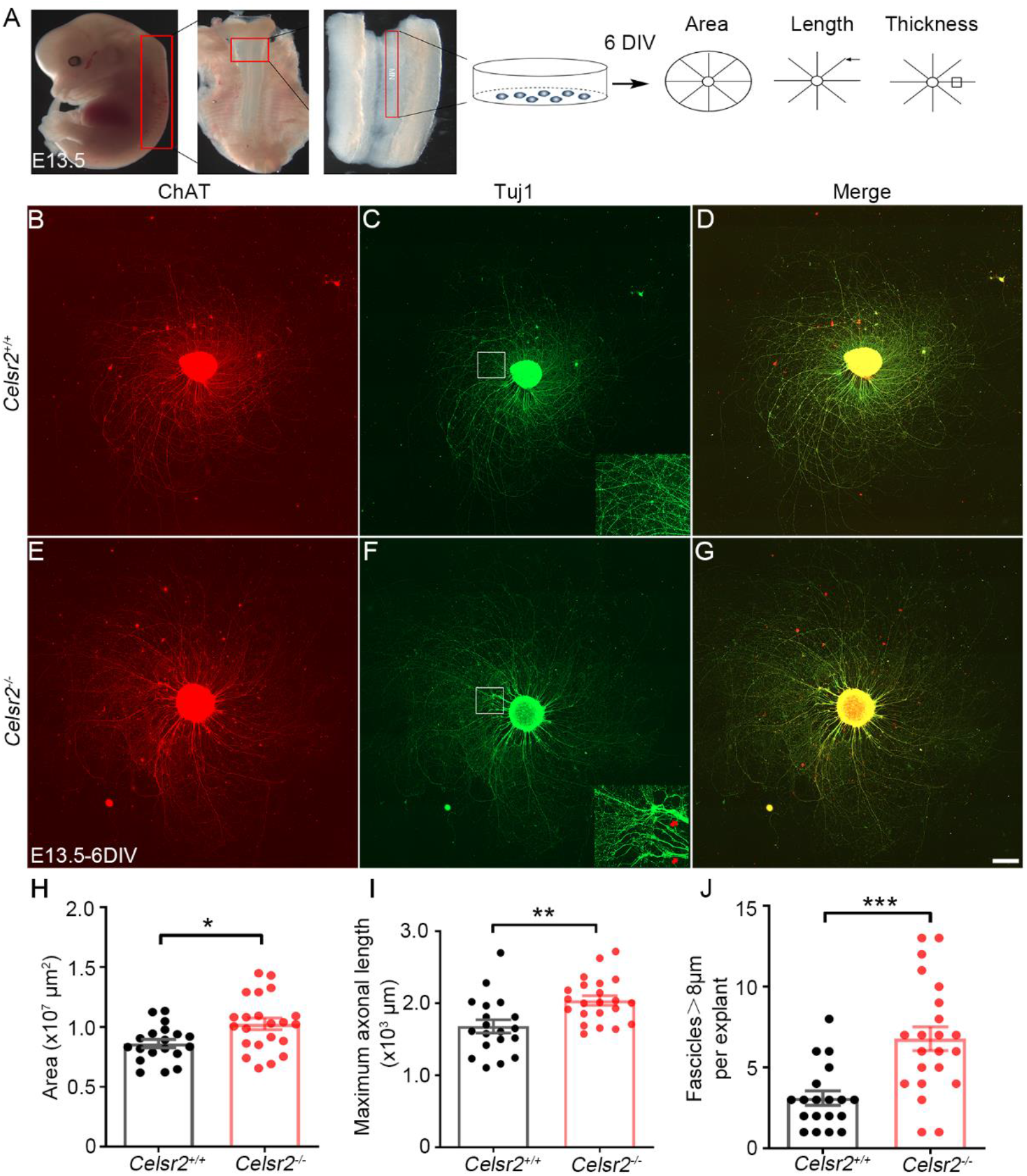
*Celsr2* knockout improves axon growth in mouse spinal motor explant culture. (A) Schema of the experimental procedure for explant culture and analysis. (B-G) Spinal motor neuron explants from E13.5 *Celsr2*^*+/+*^ (B-D) and *Celsr2*^*−/−*^ mouse embryos (E-G) were cultured for 6 DIV and then immunostained for ChAT (B, E; red) and Tuj1 (C, F; green). Both signals colocalize (D, G; yellow). Representative axonal bundles are indicated in the Insets of C and F. (H-J) Quantification of the maximal area covered by growing axons (H), maximal axon length (I), and number of large axon bundles (> 8 µm in diameter) (J). These parameters are increased significantly in the mutant compared to the control. ***, *P*<0.001; **, *P*<0.01; *, *P*<0.05; Student *t*-test. Scale bar: 200 µm in B-G.

In explant cultures, spinal motor neurons are mixed with other cells such as interneurons or glial cells that may influence motor axon growth. To check whether *Celsr2* acted in a cell-autonomous manner, we cultured isolated motor neurons from E13.5 embryos. After 6 DIV, axons were readily identified by anti-Tuj1 immunofluorescent staining (Figure 3, A and B). Like in explant cultures, the total neurite length was significantly enhanced in the mutant compared to the control (Figure 3E; 3 independent experiments, 35 neurons and 41 neurons in the mutant and control respectively; *P*<0.0001). Anti-F-actin and - Tuj1 double immunostaining labelled the growth cone and axon shaft (Figure 3, C and D). The growth cone size, estimated by F-actin immunoreactivity, was increased in mutant compared to control samples (Figure 3F; 3 independent experiments, 39 neurons and 55 neurons in the mutant and control respectively; *P*<0.0001).

**Figure 3.**
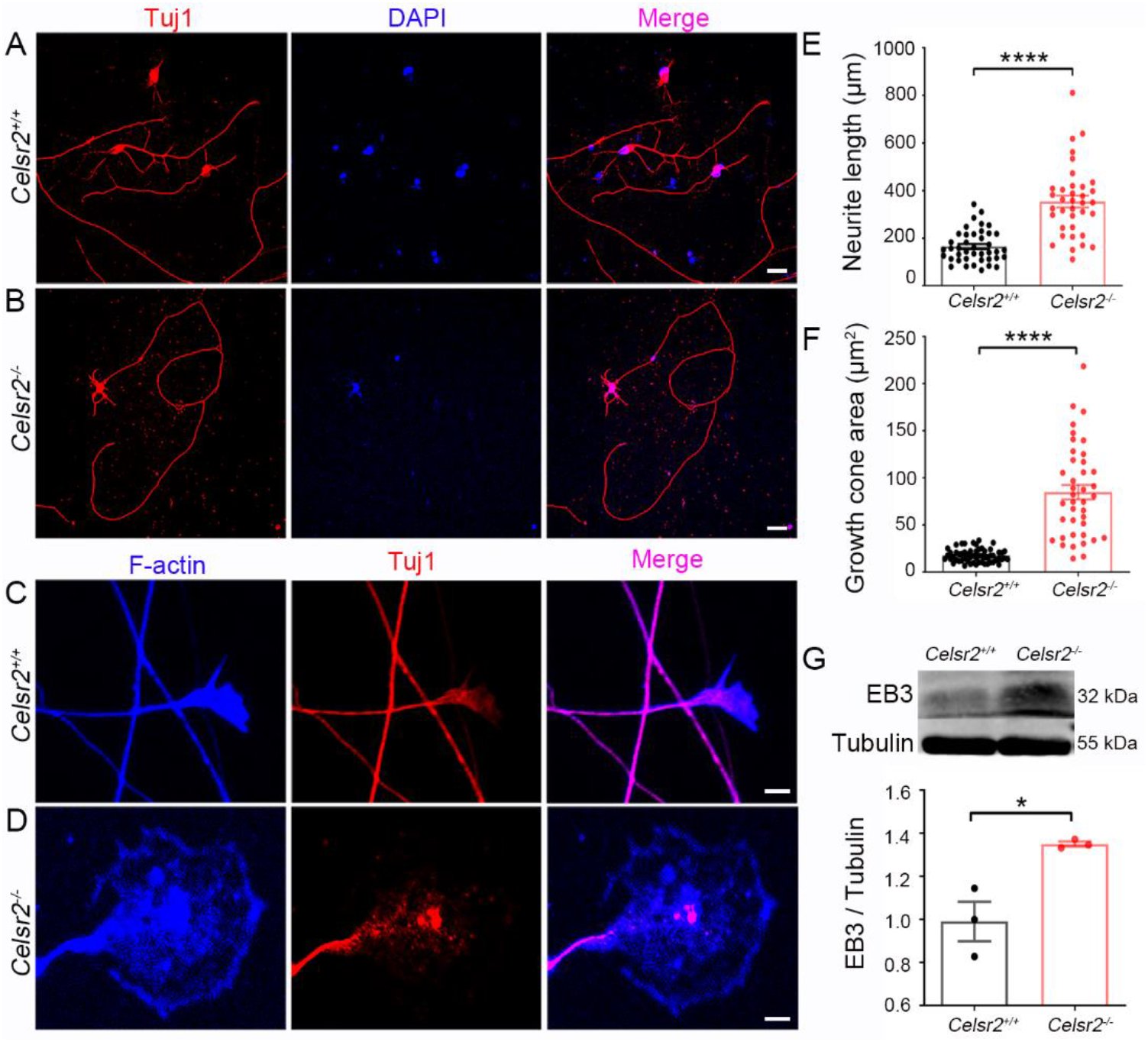
*Celsr2* knockout contributes to neurite growth in primary spinal motor neuron culture. (A, B) E13.5 spinal motor neurons from *Celsr2*^*+/+*^ (A) and *Celsr2^−/−^* (B) mouse embryos were cultured for 6 DIV and immunostained for Tuj1 (red). DAPI counterstained nuclei (blue). (C, D) Double immunostaining of cultured neurons for F-actin (blue) and Tuj1 (red) disclosed the axon shafts and growth cones. (E, F) Statistic analysis of total neurite length (E) and growth cone areas (F). (G) Protein extracts from E13.5 ventral horns of cervical spinal segments was subjected to western blots using anti-EB3 and β-III tubulin (tubulin). There was a dramatic increase of EB3 in the mutant compared to the control. *, *P*<0.05; ****, *P*<0.0001; Student *t*-test. Scale bars: 20 µm in A and B, 10 µm in C and D.

Axonal growth is a dynamic process that requires remodeling of cytoskeleton. The upregulation of the microtubule plus end-binding protein 3 (EB3) occurs in parallel to axon regeneration (Geraldo, Khanzada, Parsons, Chilton, & Gordon-Weeks, 2008). We analyzed protein content of E13.5 spinal ventral horn extracts by western blots and found that the EB3 levels were markedly higher in *Celsr2^−/−^* than in controls (Figure 3G).

Intracellular calcium is an established indicator of cytoskeleton reorganization during axon growth. In 16-DIV cultured spinal motor neurons, we studied potassium-stimulated calcium influx using the calcium sensitive dye Fura-4 AM (Supplemental Figure 1A). Fluorescent intensity of cultured neurons measured before and after potassium stimulation showed a significant increase of intracellular calcium peak in *Celsr2^−/−^* compared to control mice (Supplemental Figure 1, B and C; *P*<0.05).

### Rac1/Cdc42/JNK/c-Jun signaling is upregulated in *Celsr2^−/−^*embryonic spinal cord

Small Rho family GTPases Cdc42 and Rac1 are critical modulators of cytoskeleton organization, with their GTP-bound, active form stimulating neurite growth (Samuel & Hynds, 2010). To assess whether *Celsr2* inactivation affects the expression and/or activity of Cdc42 and Rac1, we extracted proteins from E13.5 spinal segments and pulled down GTP-bound Rac1 and Cdc42 using agarose beads conjugated to the protein binding domain of PAK1 fused to GST, followed by elution and western blot analysis with anti-Rac1 and anti-Cdc42 antibodies. Whereas total Cdc42 and Rac1 levels were similar in *Celsr2^−/−^* and control extracts, the level of active Cdc42 and Rac1 proteins was enhanced, by 1.55 (Cdc42) and 1.74 (Rac1) fold in *Celsr2^−/−^* compared to control samples (Figure 4, A, B, E and F; 3 independent experiments). As JNK and c-Jun are downstream partners regulated by Rac1 and Cdc42 during cytoskeleton reorganization (Hall, 2005), we estimated their expression using western blots (Figure 4, C and D). There was a significant increase of JNK and c-Jun concentrations in *Celsr2* mutant *versus* control samples (Figure 4, G and H; 3 independent experiments; *P*<0.05). These results suggest strongly that *Celsr2* knockdown enhances Cdc42/Rac1/JNK/c-Jun signaling in the developing spinal cord.

**Figure 4.**
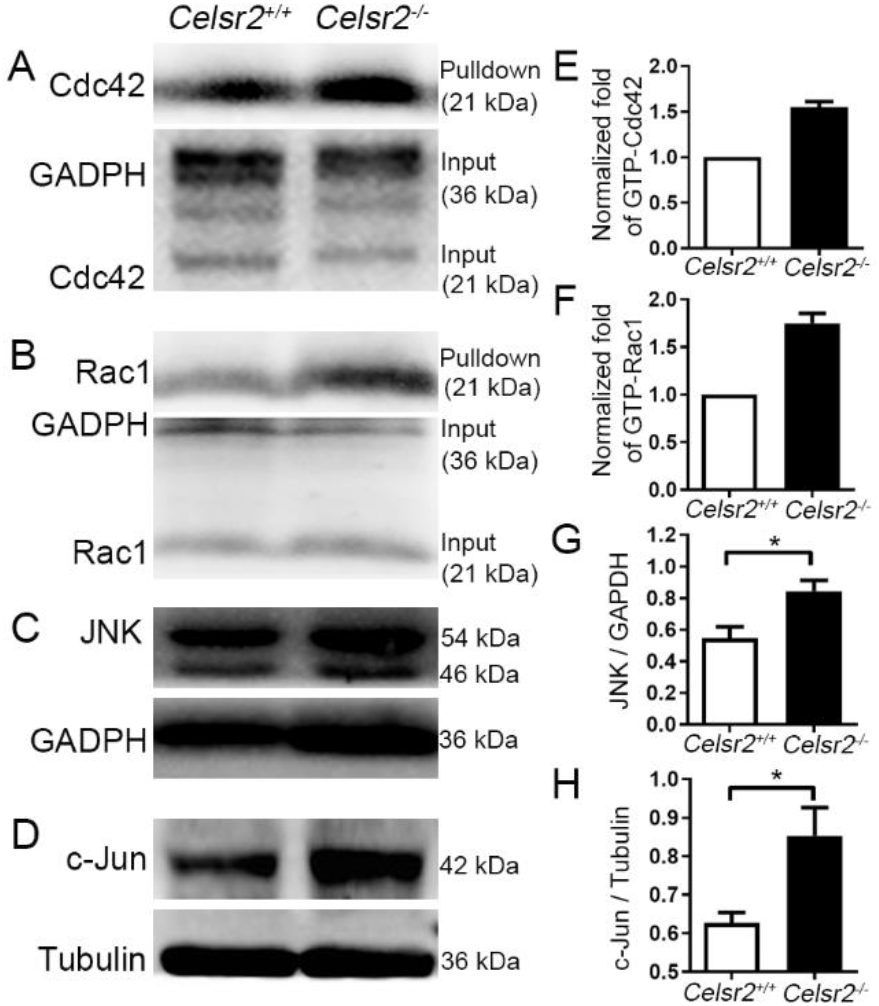
High Cdc42/Rac1 and JNK/c-Jun signaling is observed in *Celsr2^−/−^*embryonic spinal cords. (A-D) Western blots of GST-pulldown proteins and extracts from ventral columns of E13.5 cervical spinal segments were analyzed using anti-Cdc42 (A), -Rac1 (B), -JNK (C), -c-Jun (D) antibodies. Anti-GADPH or -β-III tubulin (tubulin) antibodies were used as references. (E-H) In the mutant, levels of GTP-bound Cdc42 (E) and Rac1 (F) were 1.55 and 1.74-fold of those in the control. There was a significant increase of JNK (G) and c-Jun (H) concentrations in the mutant compared to the control. *, *P*<0.05; Student *t*-test.

### *CELSR2* knockdown stimulates the growth of spinal motor axons and enhances Cdc42/Rac1 signaling in humans

To test whether CELSR2 has similar effects in humans as in mice, we used lentiviral vectors encoding CELSR2 shRNA, and evaluated their efficacy by transfecting cultured motor neurons from WPC7 human embryos. Three different CELSR2-shRNA were compared (Supplemental Figure 2) and the most efficient one was selected for study. Three days after transfection, CELSR2 mRNA level was 28% of the control (using RT-qPCR, Supplemental Figure 2A) and the CELSR2 protein was 23% of the control (western blots, Supplemental Figure 2, B and C). We cultured spinal explants from WPC7-8 human embryos and transfected explants with CELSR2 shRNA-lentivirus or scrambled shRNA-lentivirus as control. After 5 DIV, cultured explants were immunostained with anti-ChAT and -Tuj1 antibodies. In both groups, lentivirus-tagged GFP labelled cells in the explants, and some labeled neurons migrated out of the explants (Supplemental Figure 3, A and E). ChAT- and Tuj1-immunoreactivity were consistently colocalized in explants and outgrowing axons (Supplemental Figure 3, B-D and F-H), indicating that growing axons stemmed from motor neurons. Compared to mouse, the human explants developed faster, and the area covered by growing axons was much larger after 5 DIV (Figure 5, A-D) than that in mouse after 6 DIV (Figure 2, B and E). Intriguingly, in the CELSR2-shRNA group, growing axons often gathered together to form circles (11/27; Figure 5, C and C’), and some aggregated into large bundles (Figure 5, D and D’). We measured the maximal covering area and axonal length in anti-Tuj1 immunostaining preparations. Statistical analysis confirmed a significant increase in both parameters in CELSR2*-*shRNA transfected explants compared to those transduced by CELSR2 scrambled shRNA (Figure 5, E and F; 3 human embryos; 24 and 23 explants in the CELSR2*-*shRNA and the control respectively; *P*<0.0001). In addition, larger axonal bundles (>8 µm in diameter) were present after CELSR2*-*shRNA *versus* control transfection (Figure 5G; *P*<0.0001).

**Figure 5.**
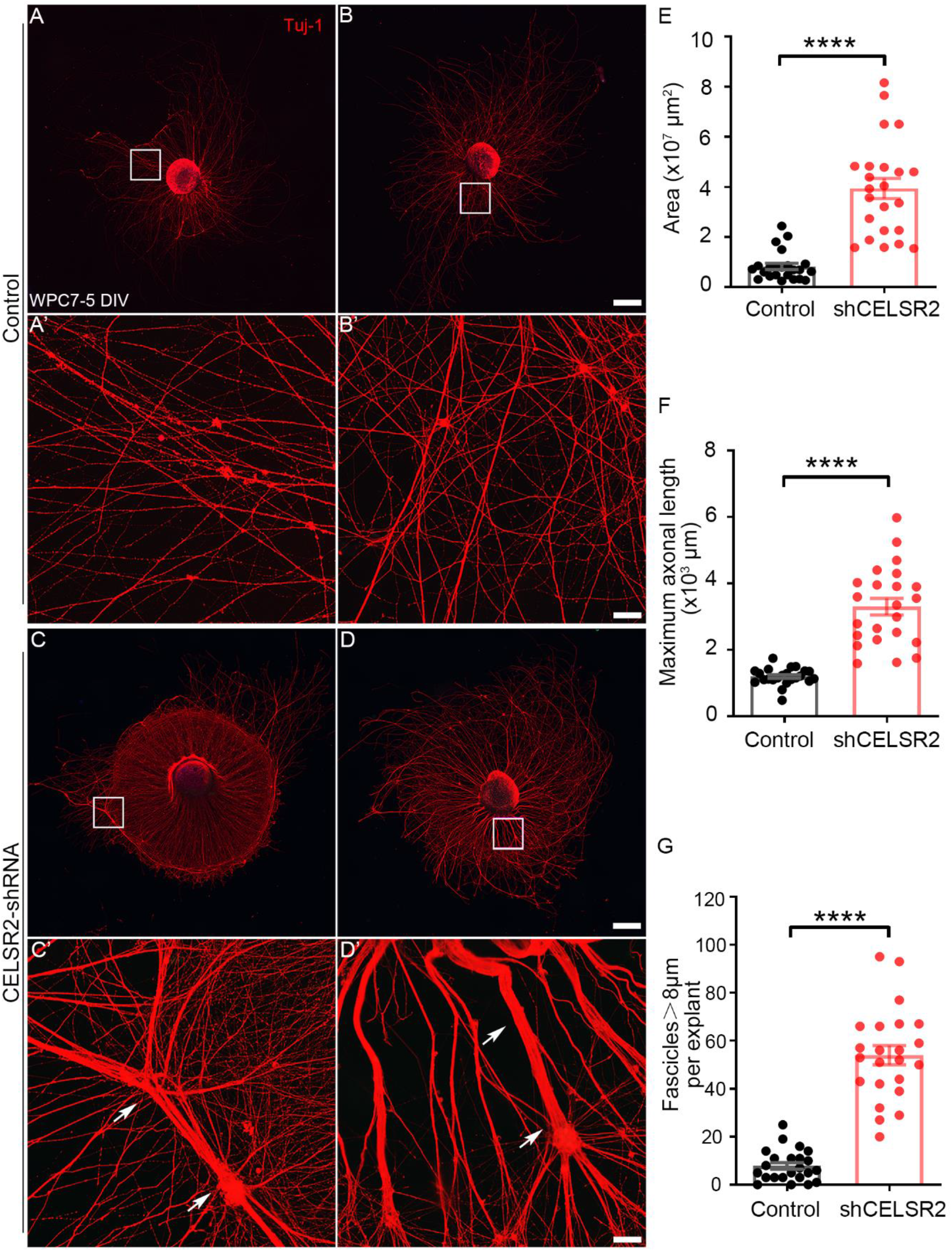
*CELSR2* knockdown increases axonal regeneration in human spinal motor explant culture. (A-D) Cultured spinal motor neuron explants from WPC7 human embryos were transfected with a CELSR2 scrambled shRNA as control (A, B) and with CELSR2-shRNA (C, D). After 5 DIV, cultured explants were immunostained for Tuj1. A’, B’, C’ and D’ are enlarged areas from A, B, C and D respectively. Upon CELSR2-shRNA knockdown, growing axons grew in circles (one example indicated in C), and formed large axonal bundles (arrows in C’ and D’). (E-G) The maximal area (E), maximal axon length (F), and the number of large axon bundles (G) were significantly increased in the CELSR2-shRNA knockdown explants compared to control. ****, *P*<0.0001; Student *t*-test. Scale bars: 500 µm in A and B, 50 µm in A’ and B’, 500 µm in C and D, 50 µm in C’ and D’.

In human primary motor neuron culture, lentivirus-tagged GFP labeled neuronal soma and neurites exactly like Tuj1-immunostaining (Figure 6, A and B). Total neurite length was increased in CELSR2-shRNA transfected neurons at 5 DIV (Figure 6C; 2 WPC7 and 1 WPC8 human embryos, 43 and 36 neuorns from the control and the CELSR2-shRNA transfection respectively; *P*<0.0001). GFP also filled growth cones which were visualized by anti-F-actin immunostaining (Figure 6, D and E). The growth cone area was considerably larger in the CELSR2-shRNA than the control group (Figure 6F; *P*<0.0001). In 5-DIV CELSR2-shRNA transfected neurons, EB3 protein levels were significantly higher than in controls (Figure 6G; 3 independent experiments; *P*<0.05). In 19-DIV cultured human motor neurons, the peak of potassium-induced intracellular calcium was more elevated in CELSR2-shRNA transfected neurons than in controls (Figure 6H; *P*<0.05).

**Figure 6.**
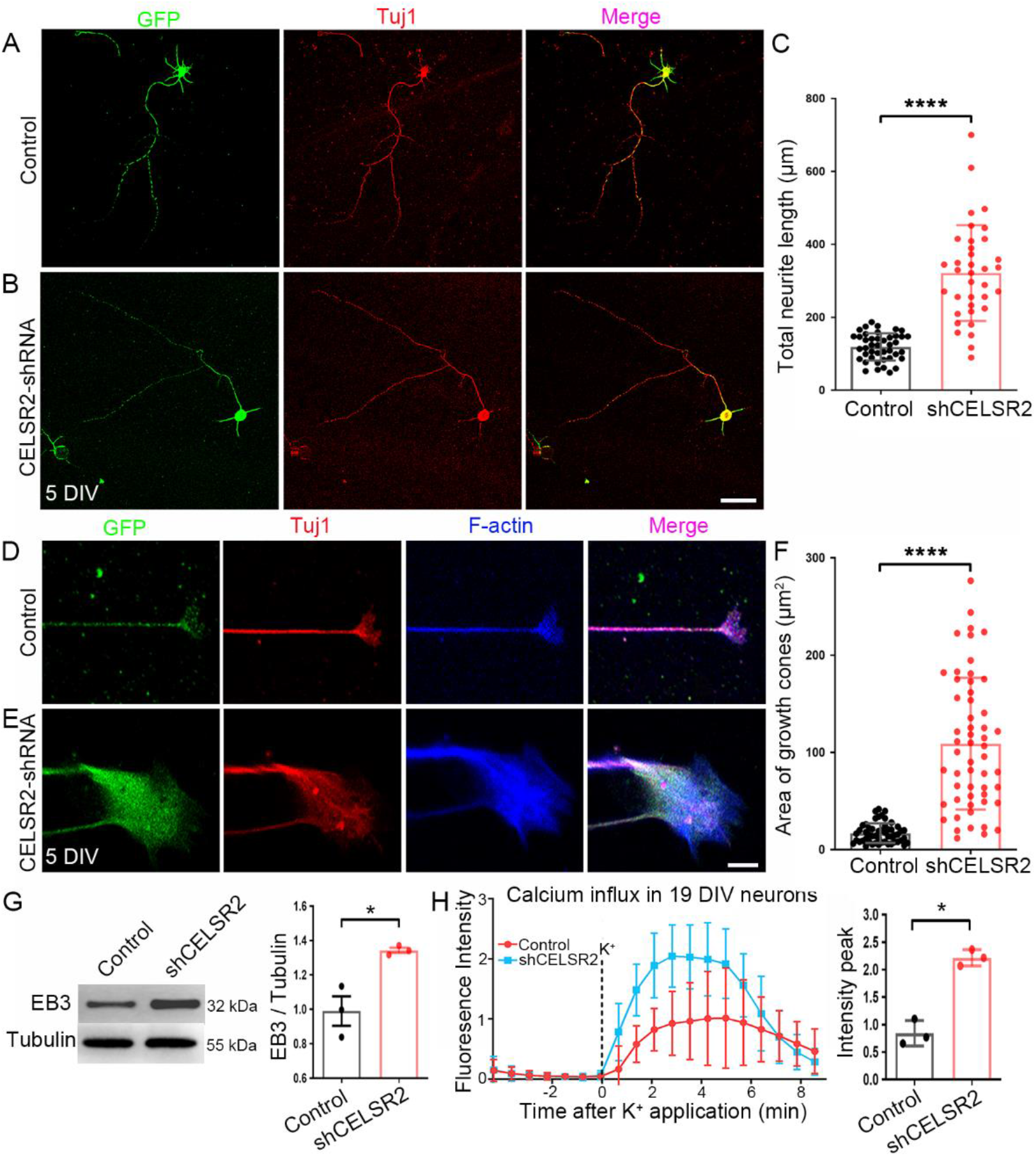
*CELSR2* knockdown promotes axonal growth in primary human spinal motor neuron culture. (A-C) CELSR2 scrambled shRNA (A, Control) and CELSR2-shRNA (B) were used to transfected cultured primary spinal motor neurons from WPC7 and WPC8 human embryos. Transfected neurons were visualized by virus-encoded GFP (green). After 5 DIV, neurons were immunostained for Tuj1 (Red). GFP and Tuj1-immunoreactivity overlapped in the somas and neurites as shown in the merged images. A significant increase of total neurite length was observed in CELSR2-shRNA transfected neurons (C). (D-F) Double immunostaining for F-actin (blue) and Tuj1 (red) reveal axon shafts and growth cones in control (D) and CELSR2-shRNA (E) transfected neurons (GFP labeling, Green). The growth cone area was significantly increased in CELSR2-shRNA *versus* control transfected neurons (F). Western blot analysis of 5-DIV cultured neurons with antibodies to EB3 and β-III tubulin (tubulin, reference), showed an increase of EB3 levels in CELSR2-shRNA transfected neurons. Intracellular calcium influx was evaluated in 19 DIV-cultured neurons by measuring Fluoro-4 AM fluorescence intensity. The curves were drawn from captured images before and after potassium application. There was an increase of the fluorescent peaks in CELSR2-shRNA transfected neurons. *, *P*<0.05; ****, *P*<0.0001; Student *t*-test. Scale bars: 50 µm in A and B, 10 µm in D and E.

In cultured motor neurons, CELSR2-shRNA transfection remarkably resulted in 1.36- and 2.08-fold increase of GTP-Cdc42 and -Rac1 proteins, respectively, compared to control, whereas total protein levels were comparable (Figure 7, A, B, E and F; proteins from 3 independent culturing experiments of 3 human embryos). The levels of JNK and c-Jun were also increased upon *CELSR2* knockdown (Figure 7, C, D, G and H).

**Figure 7.**
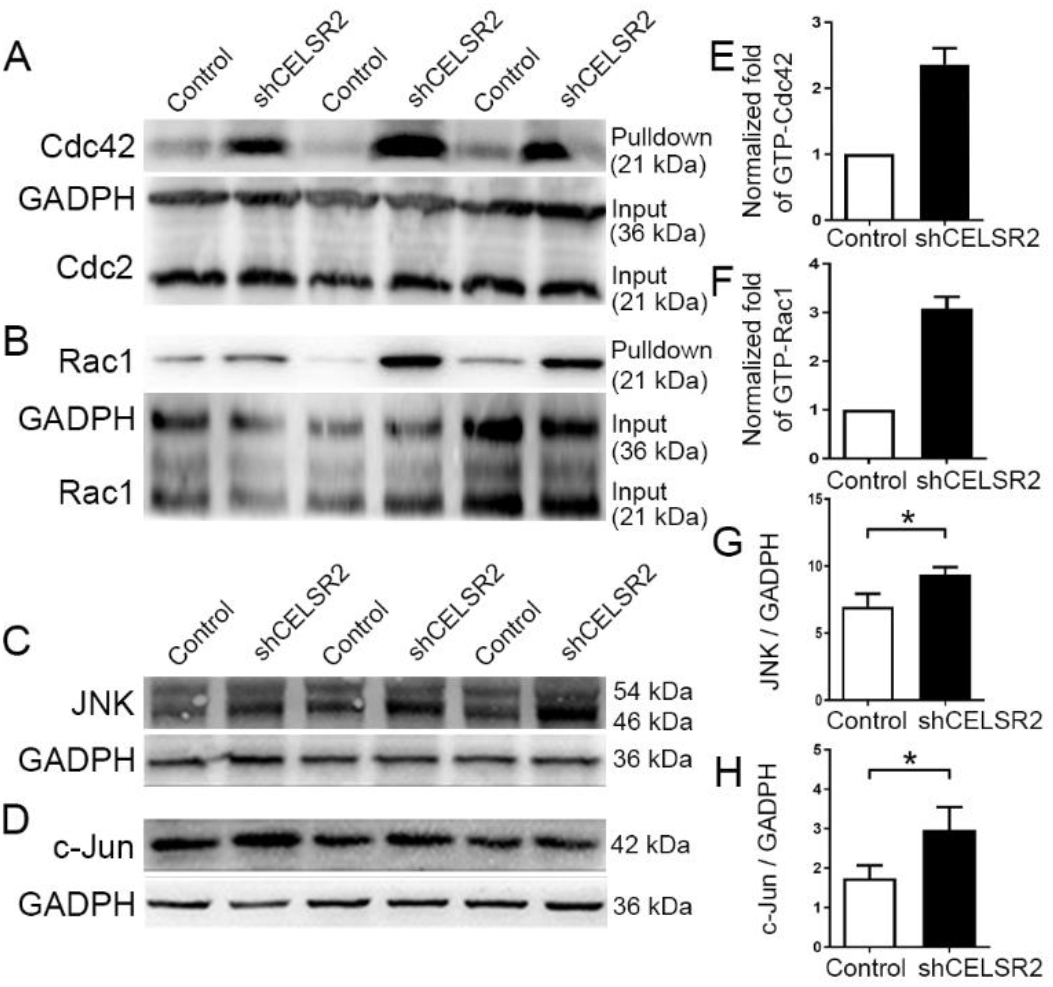
*CELSR2* knockdown enhances Cdc42/Rac1 and JNK/c-Jun signaling in cultured human primary spinal motor neurons. (A-D) After 5 DIV, proteins extracted from cultured human spinal motor neurons followed were analyzed by GST-pull down or direct western blots. Analysis with antibodies to Cdc42 (A) and Rac1 (B) showed a remarkable increase of GTP-bound Cdc42 and Rac1 in CELSR2-shRNA *versus* control transfected neurons. Concentrations of JNK (C) and c-Jun (D) were also increased in mutant samples. Anti-GADPH antibody was used as the reference. (E-H) In CELSR2-shRNA transfected neurons, the protein levels of GTP-bound Cdc42 (E) and Rac1 (F) were respectively 2.36 and 3.08 folds of those in controls. JNK and c-Jun protein levels were also significantly increased in CELSR2-shRNA transfected neurons. *, *P*<0.05; Student *t*-test.

### Inactivating *Celsr2* in motor neurons promotes axonal regeneration and functional recovery after root injury *in vivo*

To test whether *Celsr2* influences regeneration of adult motor axons *in vivo*, we used *Isl1-Cre*; *Celsr2^f/−^* mice with conditional inactivation of *Celsr2* in motor neurons. These animals develop to adulthood unremarkably, without apparent neurological deficit. We carried out brachial plexus injury and motor root re-implantation as described (Ding et al., 2014). In this model, upon reimplantation of root C6, regenerating axons can reinnervate the biceps brachii and restore a partial function (Fournier, Mercier, & Menei, 2005) that can be evaluated by monitoring elbow flexion in grooming and climbing tests. Both tests indicated a significantly improved recovery of injured right forelimb function in *Isl1-Cre*; *Celsr2^f/−^* mutants than in littermate controls (*Celsr2^f/−^*) (Figure 8, A and B). Inactivation of *Celsr2* in motor neurons decreased biceps atrophy, as evidenced by increased muscle wet weight (Figure 8, C and D), fostered biceps reinnervation with formation of new Neuromuscular junctions (NMJs), as indicated by more numerous NMJs (Figure 8E; 213 ± 16.6 *versus* 56 ± 3.5 NMJs / muscle in mutant and control respectively; n=3 animals in each group; *P*<0.001). On day 56 post-surgery, EMG recording of biceps showed a higher peak-peak amplitude value on the injured side, and an increase of the peak ratio (injured to intact sides) in the mutant compared to the control, whereas, on intact side, there were no differences of EMG amplitude and latencies between two groups (Figure 8, F-H).

**Figure 8.**
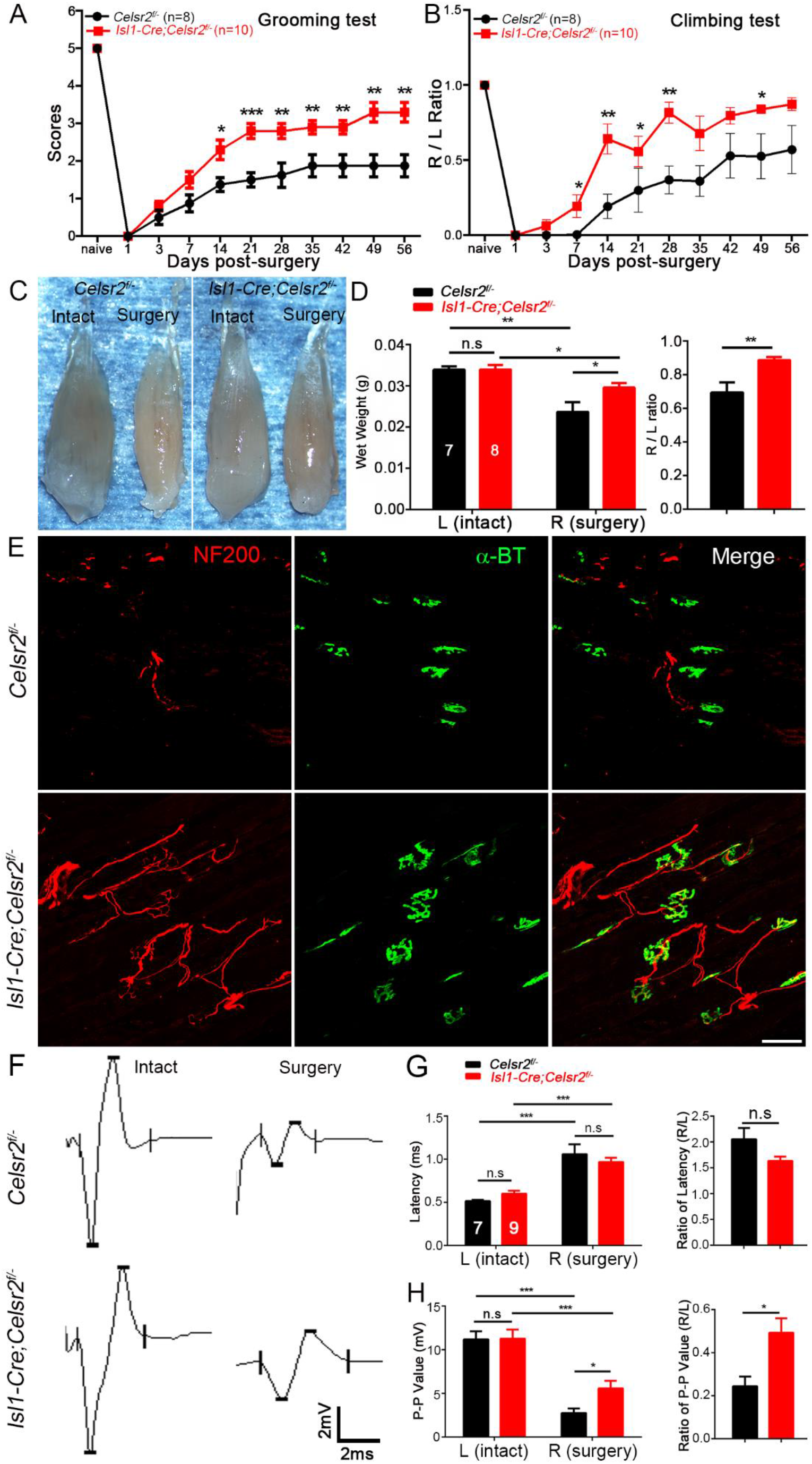
*Celsr2* conditional inactivation in spinal motor neurons improves functional recovery and NMJ formation after BPI. (A, B) After BPI, the function of the affected forelimb was assessed using the grooming (A) and climbing test (B). Scores were significantly higher in the *Isl1-Cre*; *Celsr2^f/−^* compared to littermate controls at day 14, 21, 28, 35, 42, 49 and 56 post-injury. During the climbing test, usage of injured (R) and intact (L) forelimbs was compared; the R / L ratio was increased in the *Isl1-Cre*; *Celsr2^f/−^*. (C, D) Biceps muscles collected 56 days after injury, were clearly more atrophic in the injured than in the intact side in both mutant and control mice (C). Muscle wet weight on the intact side was comparable in both two groups, whereas muscle weight on the injured side was higher in mutant *versus* controls, as reflected by the increased the R/L ratio (injured side to intact side) (D). (E) Neuromuscular junctions (NMJs) were examined using anti-NF200 and -*a*-BT double staining 56 days post-surgery. There were more numerous growing axons and NMJs in the mutant than in the control. (F-H) EMG of biceps were recorded 56 days post-surgery (F). The latencies were increased after injury, with no difference between both groups (G). Denervation resulted in a significant decrease of the peak-peak amplitude in both groups, but the amplitudes and the ratio of the injured to the intact muscle was significantly higher in the mutant compared to the control (H). *, *P*<0.05; **, *P*<0.01; ***, *P*<0.001; Student’s *t*-test. Scale bar: 50 µm in E.

In *Isl1-Cre*; *Celsr2^f/−^* mice, ChAT immunolabeling of C6 spinal segments showed that more spinal motor neurons survived after root ablation/reimplantation than in control mice in terms of increased ChAT positive cells and higher operated/intact ratio (Figure 9, A and B). Similarly, in semithin sections of musculocutaneous nerves, an increase of total axon numbers on injured sides and of the ratio of injured to intact sides was found in the mutant compared to the control (Figure 9, C-E).

**Figure 9.**
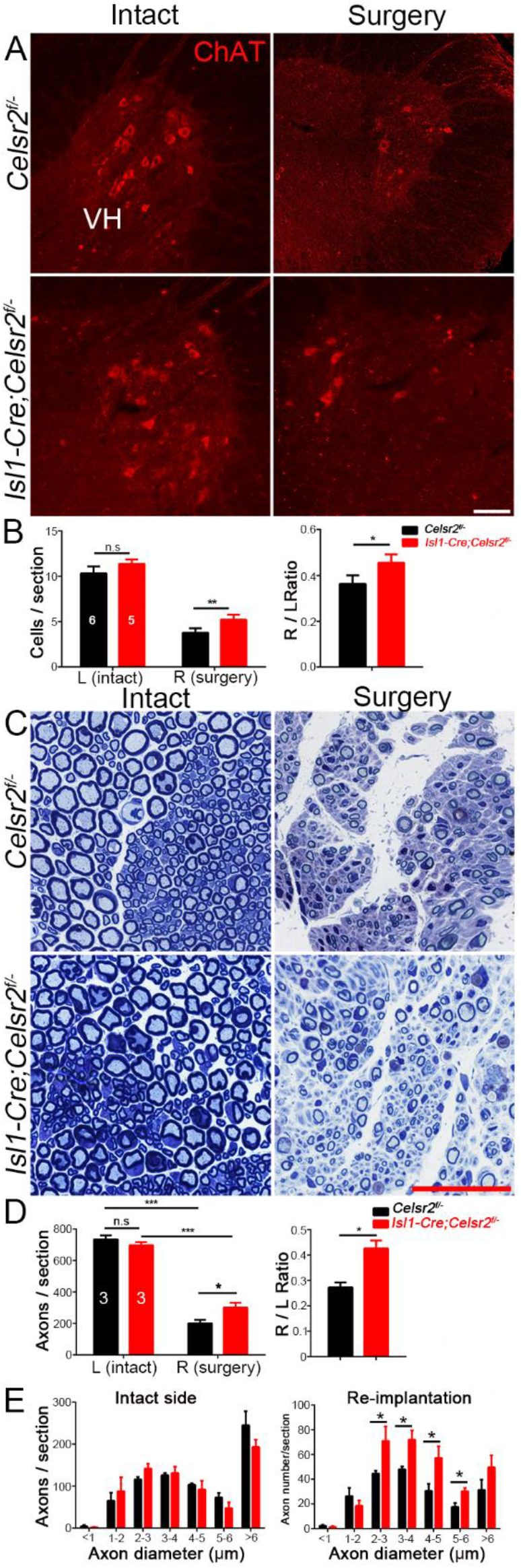
Improved spinal motoneuron survival and axon regeneration in *Celsr2* conditional knockout mice after BPI. (A, B) C5-C7 spinal sections immunostained for ChAT, showed a decrease of spinal motor neurons in the ventral horn (VH) on the operated side 56 days after injury (A). Numbers of ChAT-positive neurons on the intact side were comparable in two groups, but higher in the mutant than in control animals on the injured side, as illustrated by the increased ratio of the injured (R) to the intact (L) sides in mutant *versus* control (B). (C-E) Toluidine blue staining of musculocutaneous nerves after surgery (C) showed that axon number was comparable on the intact side in two genotypes, but higher in the mutant on the injured side (D). The ratio of the injured to the intact side (R/L) was higher in the mutant (D). The distribution of axon according to their diameter showed no differences in two genotypes on the intact side, and an increased number of axons with 2-6 µm diameters in operated side in mutant relative to control (E). *, *P*<0.05; **, *P*<0.01; ***, *P*<0.001; n.s, not significant; Student’s *t*-test. Scale bars: 100 µm in A, 50 µm in C.

To evaluate Cdc42/Rac1-JNK/c-Jun signaling after root lesion, we collected spinal samples from C5-C7 ventral columns 3 days after surgery for analysis with western blots (Figure 10, A-D), and observed results consistent with those obtained in culture conditions. In the mutant, levels of GTP-bound Cdc42 and Rac1 were about 2.2 fold and 1.9 fold of those in controls (Figure 10, E and F), and there was a significant increase of JNK and c-Jun protein levels (Figure 10, G and H; 6 animals in each group, samples from 2 animals were mixed together in the same group, 3 independent experiments).

**Figure 10.**
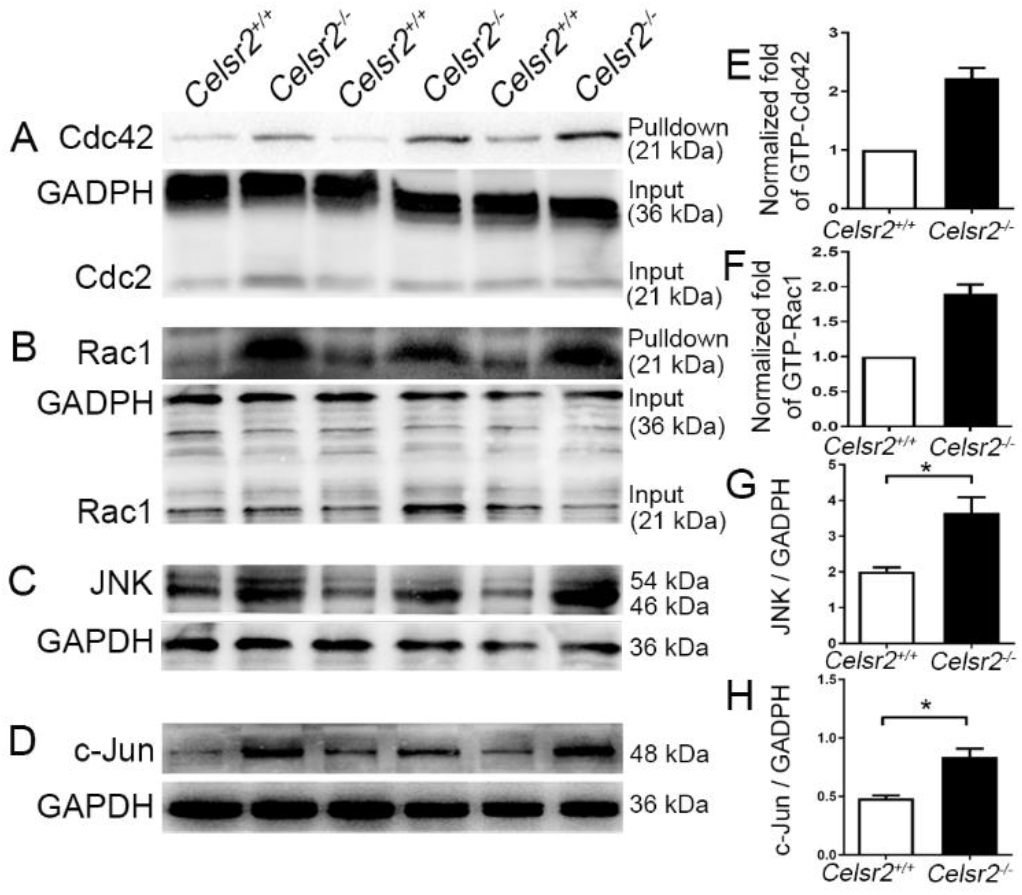
*Celsr2* knockout increases Cdc42/Rac1 and JNK/c-Jun signaling in injured ventral horns. (A-D) Western blots analysis of GST-pulldown proteins and samples from spinal ventral columns 3 days after BPI, using anti-Cdc42 (A), -Rac1 (B), -JNK (C), -c-Jun (D) antibodies. Anti-GADPH antibody was used as references. (E-H) GTP-bound Cdc42 (E) and Rac1 (F) proteins, as well as concentrations of JNK (G) and c-Jun (H) were increased in mutant compared to control samples. *, *P*<0.05; Student *t*-test.

## Discussion

Here, we report that downregulation of *Celsr2* is instrumental to axon regeneration and functional recovery of injured motor neurons. First, we showed that Celsr2 is highly expressed in spinal motor neurons. Second, we assessed neurite growth in primary cultures, and axon outgrowth in explant cultures from mouse and human embryonic ventral spinal cord, and found that motor axon growth was drastically increased upon *Celsr2* inactivation (mouse) or knockdown (human). Third, following C5-C7 root avulsion and C6 motor root re-implantation, motor neuron survival and motor axon regeneration were improved in adult mice with *Celsr2* inactivation, compared to control animals. Finally, *Celsr2* inactivation correlated with upregulation of active Rac1 and Cdc42, and of JNK, and c-Jun. Altogether, our results provide strong evidence that Celsr2 regulates negatively the regeneration of motor axons, by downregulating the Rac1/Cdc42 and JNK/c-Jun pathway.

Together with its two paralogs Celsr1 and Celsr3, Celsr2 (aka Adgrc1-3) form a small family of adhesion G-protein coupled receptors. Genetic studies in mice showed that Celsr2 and Celsr3 act synergistically during neural development, particularly in regulating axon guidance and ependymal cilia biogenesis (Tissir & Goffinet, 2013). In most cases, inactivation of *Celsr2* causes milder embryonic phenotypes than that of *Celsr3*, while joint inactivation generates severe phenotypes that mimic closely those resulting from inactivation of *Fzd3*, a member of the frizzled family of Wnt receptors (Hua et al., 2013). Celsr2 and Celsr3 can co-immunoprecipitate with Fzd3 (Chai et al., 2014), and this might explain similarity of mutant phenotypes. Neural specific inactivation of *Rac1* impairs extension of motor axons of the peroneal nerve (that innervate dorsal muscles of the hindlimb) (Hua et al., 2015), a phenotype identical to those observed when *Fzd3* or *Celsr2 &3* are conditionally inactivated in motor neurons. Given similarities of mutant phenotypes, our findings that Celsr2 regulates Rac1/Cdc42 and JNK signaling negatively, rather than positively, were unexpected. A possible explanation is that the Wnt/PCP signaling might stimulate the Rac1/Cdc42 pathway during development, while inhibiting the same pathway during axon regeneration. In support of this, at least two examples of negative regulation of the Rac1/Cdc42-JNK pathway by Wnt/PCP signaling have been published, namely the effect of Fzd6, the closest paralog of Fzd3 (Abidin, Owusu Kwarteng, & Heinonen, 2015), and of Fzd8 working together with Celsr2 in hematopoietic stem cells (Sugimura et al., 2012). The Celsr2 ectodomain is implicated in homophilic interactions (Shima et al., 2007), and could mediate repulsion between axons, accounting for increased fasciculation upon Celsr2 inactivation. In line with this, in *Drosophila*, Flamingo mediates homophilic repulsion of homologous neurons at the dorsal midline, and mutations result in formation of neurite tangles and dendrites overgrowth (Gao, Kohwi, Brenman, Jan, & Jan, 2000).

Previous studies outlined a role of JNK in PCP and noncanonical Wnt signaling, implicating, among others, the Wnt5a ligand and frizzled receptor Fzd6, a close relative of Fzd3 (Cheyette et al., 2002; Lutze et al., 2019; Yamanaka et al., 2002; Yang & Mlodzik, 2015). Our data are a first indication that Celsr2 acts via the same pathway. Mechanistic links between frizzled and Rac1/Cdc42 remain incompletely understood but could involve Dishevelled adaptors and formins such as Daam1-2 (Habas, Dawid, & He, 2003). The Rac1/Cdc42 -JNK/ c-Jun signaling module is better documented. Cdc42 and Rac1 are two important small GTPases of the Rho family, and their active form regulates actin dynamics in navigating growth cones, thereby promoting axon growth (Samuel & Hynds, 2010). Downstream of Rac1/Cdc42, the pathway sequentially involves MAP3K mixed lineage kinases such as DLK, as well as MKK7/4 and JNK1-3 (Jin & Zheng, 2019; Schellino, Boido, & Vercelli, 2019). It is coordinated by docking adaptors such as JIP1/POSH (Kukekov, Xu, & Greene, 2006; Nihalani, Meyer, Pajni, & Holzman, 2001; Xu, Kukekov, & Greene, 2003). The Rac1/Cdc42 – JNK/c-Jun pathway is a “double edged sword”, with variable, often paradoxical actions. On the one hand, it can mediate neuronal apoptosis, for example upon nerve growth factor deprivation (Xu, Maroney, Dobrzanski, Kukekov, & Greene, 2001) and is implicated in Wallerian axon degeneration (Miller et al., 2009). On the other hand, upregulation of c-Jun is correlated with axon regeneration after injury (Raivich et al., 2004). JNK is necessary for the nuclear translocation of c-Jun to maintain cytoskeletal integrity during axon extension (Chang, Jones, Ellisman, Goldstein, & Karin, 2003), and DLK activation is required for axon regeneration (Nix & Bastiani, 2012).

Besides its regulation of Rac1/Cdc42 signaling, *Celsr2* inactivation results in an increase of the microtubule plus end binding protein EB3 and stimulates potassium-induced calcium influx. EB3 interacts not only with microtubules, but also with actin filaments, via drebrin, and those interactions promote growth cone formation and neurite extension (Geraldo et al., 2008). Similarly, an influx of calcium could activate calcium/calmodulin-dependent protein kinase II, which regulates F-actin to stimulate growth cone during development and after injuries (Xi et al., 2019).

In conclusion, our data uncover a previously unknown role of Celsr2 in the regulation of axon regeneration, via Rac1/Cdc42 and possibly other mechanisms. The fact that concordant data were observed in human and mice, and in both embryonic and adult axons, suggests strongly that this function of Celsr2 is physiopathologically relevant, and that Celsr2 is a potential target to promote neural repair.

## Materials and methods

### Animals

Animal procedures were performed following recommendations in the Guide for the Care and Use of Laboratory Animals of the National Institutes of Health. The protocol was approved by the Laboratory Animal Ethics Committee at Jinan University (Permit Number: 20111008001). The generation of *Celsr2*^-/-^ and *Celsr2*^*LacZ*^ mice was described previously (Tissir et al., 2010). *Isl1-Cre*; *Celsr2^f/−^* conditional knockout animals were obtained by crossing *Celsr2*^-/-^ with *Isl1-Cre* mice to generate *Isl1-Cre;Celsr2*^*+/-*^ mice which were mated with *Celsr2*^*f/f*^ mice. To stage embryos, animals were mated in the afternoon and vaginal plugs were checked on the next morning, noted as embryonic day 0.5 (E0.5).

### Human embryonic samples

Human embryos were collected following drug-induced abortions at Guangzhou Women and Children’s Medical Center. All procedures were approved by the Medical Ethics Committees of the hospital (ref file 2016041303) and in agreement with the Helsinki convention. Informed consent was obtained from both parents, and pregnant women had no reported disease history. Embryos were transferred into culture medium (DMEM-F12 plus 50 U/ml penicillin-streptomycin) immediately after expulsion. The age was estimated by the date of the mother’s last menstruation, and by measurement of Crown Rump length, according to a growth chart (Hern, 1984).

### Motor neuron explant culture and analysis of axon growth

Mouse embryonic motoneuron explant culture was performed as described (Wang & Marquardt, 2012). Briefly, cervical spinal motoneuron explants from E13.5 *Celsr2*^*+/+*^ (control) and *Celsr2^−/−^* embryos were minced and cultured on coverslips coated with poly lysine (0.75μg/cm^2^, BD, Biocoat) and laminin (0.9 μg /cm^2^), in 24-well plates (Corning). Plates were kept at room temperature for 30 min to allow explants to attach, and then transferred to a cell culture incubator. The culture medium consisted in neurobasal medium (Invitrogen) supplemented with 2% (vol/vol) B27 (Invitrogen), 50 U/ml penicillin-streptomycin, 2 mM L-glutamine and 20 ng/ml BDNF (R&D Systems). In each group (control and *Celsr2* mutant), three E13.5 litters of 5-7 embryos were used, and 19-22 explants were suitable for analysis. Human spinal motoneuron explant culture was performed from 7 post conceptional weeks (WPC7) embryos, in 12-well plates, using similar procedures. After 24-hour culture, 2×10^9^ transduction units (TU) / ml lentivirus encoding CELSR2-shRNA, or CELSR2 scrambled shRNA as control was added to the culture medium and replaced by fresh culture medium after 48 hours. After 6 DIV for mouse and 5 DIV for human, cultured explants were fixed and immunostained with anti-Tuj1 (1:1000; ab18207; Abcam) and - ChAT (1:500 dilution; AB144P, Millipore) antibodies. The maximal area covered by growing axons was estimated using ImageJ (Matsunaga, Hatta, Nagafuchi, & Takeichi, 1988). In each explant, 4 axes (8 directions in total) were drawn with the explant at the center; the maximal axon length was measured in each direction and the average was calculated. To evaluate fasciculation, axonal bundles with diameters larger than 8 μm were counted in the region 150-200 μm away the explant surface (Jaworski & Tessier-Lavigne, 2012).

### Primary spinal motor neuron culture and analysis

Ventral horns of cervical spinal cords from E13.5 mouse or WPC7-8 human embryos were dissected out in cold DPBS (Hanks Balanced Salt Solution without Ca^2+^ and Mg^2+^ at pH 7.4). After trypsinization (0.5% trypsin, Gibco) and cell counting, isolated cells (2.5 × 10^3^ cells/cm^2^) were seeded on poly-lysine and laminin coated coverslips, in 12-well plates, and cultured in neurobasal medium (Invitrogen) plus 2% B27, 20 ng/ml BDNF, 50 U/ml penicillin-streptomycin and 2 mM L-glutamine. Half of the medium was replaced by fresh medium every 2-3 days. To estimate the effects of *CELSR2* inactivation in human spinal neurons, lentivirus (2×10^9^ transduction units (TU)/ml) encoding CELSR2-shRNA, or CELSR2 scrambled shRNA as control was added to the culture medium after 1 DIV, and fresh culture medium was added 48 hours later. After 5 DIV, cells were fixed and stained with mouse anti-Tuj1 (1:1000; ab18207; Abcam) and TexRed-Phalloidin (1:50, Life Technologies Corporation). Neurite length and growth cone areas were measured using Imaris and Image J. Data were collected from at least 3 independent cultures, and a total of 50–60 neurons were used for quantification in each group.

### Measurement of intracellular calcium

After 16 DIV for mouse and 19 DIV for human, cultured neurons were washed 3 times with HBSS, incubated in complete medium supplemented with 5 μM Fura-4 AM (DOJINDO Chemical Technology) for 30 min, and rinsed with HBSS. Cover slips were transferred to a culture chamber under an inverted microscope (Axioplan 2; Zeiss, Germany) and perfused with calcium-imaging buffer at a rate of 2.0 ml/min for 13 min. 50 mM KCl was applied to stimulate calcium influx. Fluorescence density was analyzed to estimate the intracellular calcium peak, using ImageJ software. Three litters (6-7 embryos in each litter) of E13.5 mouse embryos and 3 human embryos (2 at WPC7 and 1 at WPC8) were used for spinal neuron culture.

### Animal surgery

Adult mice (2-3 months old, 24-28 g) were selected for surgery and anesthetized with avertin (1.25%, 20μl/g, intraperitoneal injection). C5-C7 spinal segments were exposed under an operating microscope following unilateral hemilaminectomy. Right C5-C7 dorsal and ventral roots were dissected out; 2-3 mm of spinal roots in segments of C5 and C7 were removed to prevent reconnection to spinal cord, and the right C6 ventral root was re-implanted to its initial location. During the operation, a thermostatic surgical pad was used to maintain normal body temperature. After the procedure, animals were housed in individual cages. They resumed drinking and eating within 24 hours and recovered uneventfully.

### Behavioral tests

Behavioral tests were carried out by an experimenter blind to mouse genotypes.

#### Grooming test

C6 spinal motor axons travel in musculocutaneous nerve and innervate the biceps to control elbow flexion. Functional recovery was assessed by the grooming test (Bertelli & Mira, 1993), and the scores were determined by monitoring the position that the right forelimb could reach. After water was sprayed on the animal’s head, the forelimb movements were recorded for 2 min with a video camera and the scores were given as described (Ding et al., 2014). After 24 hours of surgery, successful animals had the scores of 0, and those with the score of above 0 were excluded.

#### Climbing tests

Climbing tests evaluated the usage of forelimbs (Han et al., 2015; Starkey et al., 2005). Briefly, mice were placed in a clear Perspex cylinder (170 mm in height, 90 mm in diameter) and the times of forelimb touching the wall were counted during a video recording.

The ratio of hits by forelimbs on the operated *versus* control side was calculated.

### Electromyography (EMG)

The musculocutaneous nerve and biceps brachii were exposed under a surgical microscope, under 1.25% avertin anaesthesia. A stimulating electrode was placed on the musculocutaneous nerve and a bipolar recording electrode was inserted in the center of the biceps, with a grounding electrode in the subcutaneous tissue. Similar stimulations (30 µA, 30 ms) were applied in all animals. EMG signals were collected with a multi-channel system (VikingQuest EMG/EP System, Nicolet, USA). Individual responses were measured 6 times, at 2-min intervals, on left and right sides, and the mean from 6 trials represented one sample. The peak-to-peak amplitude of evoked potentials was measured and expressed as the ratio between the operated (R) and the unoperated (L) sides. After recording, the biceps brachii were fixed in 4% paraformaldehyde to estimate the R/L weight ratio.

### Histology and immunohistochemistry

C5-C7 spinal segments or musculocutaneous nerve were cut into 15-µm-thick frozen sections for immunostaining. The blocking buffer was composed of 5% goat serum and 3% bovine serum albumin diluted in 0.1M phosphate buffer saline (PBS). Signal was detected with Alexa fluor 546 or 488 coupled secondary antibodies (1:1000, Invitrogen). Primary antibodies were: goat anti-ChAT (1:500, ab144p, Millipore), chicken anti-*β*-gal (1:500, ab9361, Abcam) and rabbit anti-Isl1 (1:1000, Abcam). To visualize neuromuscular junction (NMJ), the biceps were collected 50 days post-surgery and the wet weight was recorded. Seventy-µm horizontal sections were prepared with the a sliding microtome (Leica, Germany) and double stained with rabbit anti-NF200 (1:500, n4142, Sigma) and α-BT (1:1000, Molecular probes, USA). To analyze axon number in musculocutaneous nerves, tissues were fixed with 2.5% glutaraldehyde plus 2% paraformaldehyde and 500 nm semi-thin sections were stained with 1% Toluidine Blue; images were taken under a 63x oil objective.

### Rac1/Cdc42 activity assay

Levels of GTP-bound Cdc42 and Rac1 proteins of E13.5 mouse samples containing cervical spinal motor neurons, 5-DIV cultured human spinal motor neurons and C5-C7 ventral horns of adult animals 3 days after surgery were estimated using a Rac1/Cdc42 Activation Assay Kit (#17-441, Merck Millipore). Briefly, samples were homogenized in the lysis buffer (125 mM HEPES (pH 7.5), 750 mM NaCl, 5% Igepal CA-630, 50 mM MgCl2, 5 mM EDTA, 10% glycerol and protease inhibitors). Lysates were incubated with glutathione-agarose beads conjugated with the PAK1 Protein Binding Domain fused to GST (GST-PBD) at 4 °C for 60 min and the beads were washed three times in lysis buffer. GTP-bound proteins were eluted in sample gel buffer and subjected to western blots.

### Western blots

Protein extracts from E13.5 mouse cervical segments, C5-C7 ventral horns of adult animals 3 days after surgery, 5-DIV cultured human motoneuron explants and proteins eluted from glutathione agarose beads were analyzed on 10% sodium dodecylsulfate polyacrylamide gels and then transferred to 0.45 μm nitrocellulose membranes. The following primary antibodies were used: anti-β-tubulin (Tuj1) rabbit polyclonal antibody (1:1,000; ab18207, Abcam), anti-GAPDH mouse polyclonal antibody (1:1,000; ab8245, Abcam), anti-JNK rabbit polyclonal antibody (1:1,000; 9252, CST), anti-c-Jun rabbit polyclonal antibody (1:1,000; 3270S, CST), anti-EB3 rat polyclonal antibody (1:1,000; ab53360, Abcam), anti-GSK-3β polyclonal antibody (1:1,000; 9315, CST), anti-Rac1 mouse polyclonal antibody (1:1,000; 610650, BD Pharmingen), anti-CDC42 rabbit polyclonal antibody (1:1,000; 11A11, CST); the secondary antibodies include Peroxidase anti-rabbit IgG (1:5,000, ab6721, Abcam) and peroxidase anti-mouse IgG (1:10,000; Vector Laboratories). Immunoreactivity was detected using an enhanced chemiluminescence (ECL) detection kit (1705061, Bio-Rad).

### Quantification and statistics

Data were presented as mean ± SEM. Results of behavioral tests were analyzed with two-way ANOVA and other comparisons were made using Student *t*-test of two-independent samples. The cell count and fluorescence density were assessed using ImageJ software.

## Author contributions

L. Zhou and Y. Qu designed research studies; Q. Wen, H. Weng, T. Liu conducted experiments, acquired and analyzed data; L. Yu, J. Qin and T. Zhao acquired data and provided reagents, F. Tissir provided reagents and analyzed data; Q. Wen, H. Weng and L. Zhou prepared the draft; L. Zhou, Y. Qu and F. Tissir corrected the manuscript.

## Acknowledgements

We wish to thank Meizhi Wang, Lei Shi for technical assistance, and Prof. Kwok-Fai So for critical comments. This work was supported by the following grants: National Natural Science Foundation of China (81971148, L. Zhou and 31671067, Y. Qu), Guangzhou Key Projects of Brain Science and Brain-Like Intelligence Technology (20200730009, L. Zhou), Guangdong grant ‘Key technologies for treatment of brain disorders’ (2018B030332001, L. Zhou), Health and Medical Collaborative Innovation Major Projects of Guangzhou (201803040016-2, L. Zhou; 201604046028, L. Zhou and KF. So), Programme of Introducing Talents of Discipline to Universities (B14036), Outstanding Scholar Program of Bioland Laboratory (Guangzhou Regenerative Medicine and Health Guangdong Laboratory; 2018GZR110102002), Guangdong Natural Science Funds for Distinguished Young Scholars (2016A030306001, Y. Qu), Guangdong Province Special Support Program (2015TQ01R837, Y. Qu), and Guangdong-Hong Kong-Macao Greater Bay Area Center for Brain Science and Brain-Inspired Intelligence Fund (NO. 2019017).

