## Supplemental Figure 1, Supplemental Figure 2, Supplemental Figure 3 for "Regeneration of spinal motor axons: Negative regulation by Celsr2 implicating Cdc42/Rac1 and JNK/c-Jun signaling"

### Supplemental Materials

**Supplemental Figure 1. *Celsr2* inactivation increases calcium influx in cultured mouse spinal motor neurons.**

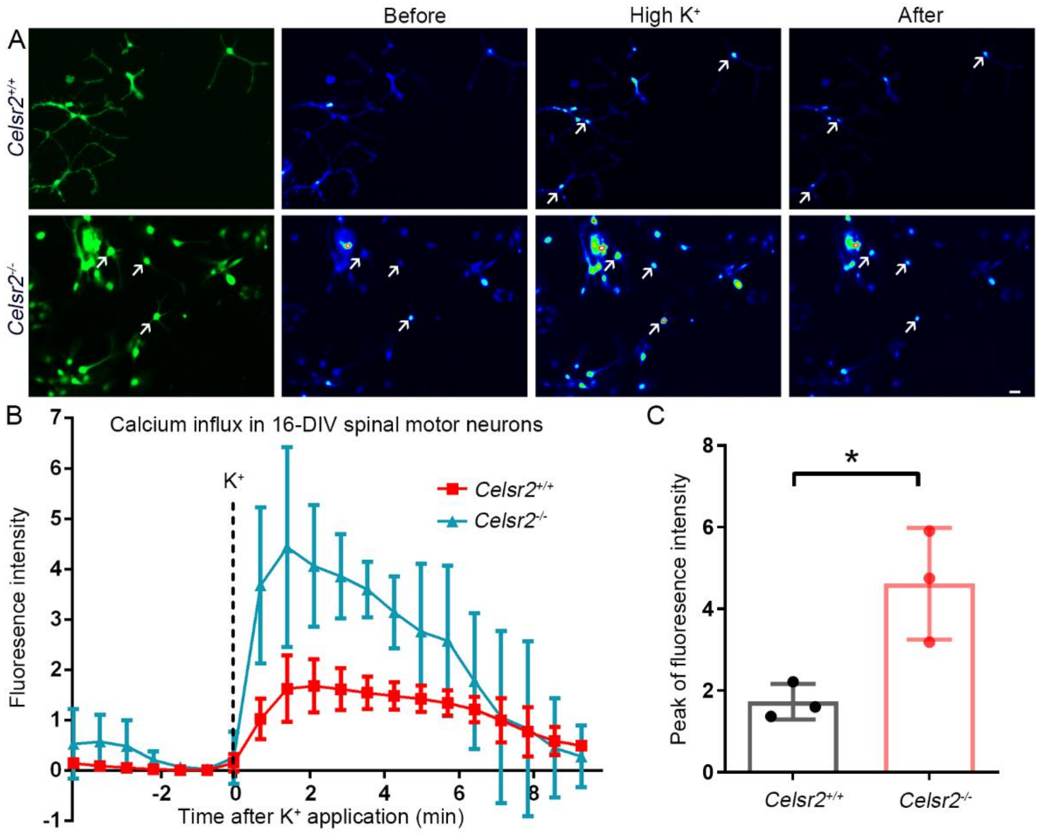


(A) After 16 DIV, cultured mouse spinal motor neurons were incubated with the calcium sensitive dye Fura-4 AM and images were captured before and after potassium stimulation, in *Celsr2*$^{+/+}$ (Control) and *Celsr2*$^{-/-}$ neurons. Representative Fura-4 AM-labelled neurons are indicated by arrows.

(B, C) Statistical analysis showed a significant increase of intracellular calcium peak in *Celsr2*$^{-/-}$ compared to control cells.

*, $P<0.05$; Student *t*-test. Scale bar: 20 µm

**Supplemental Figure 2.** ***CELSR2*-shRNA efficiently knocks down CELSR2 expression in cultured human spinal motor neurons.**

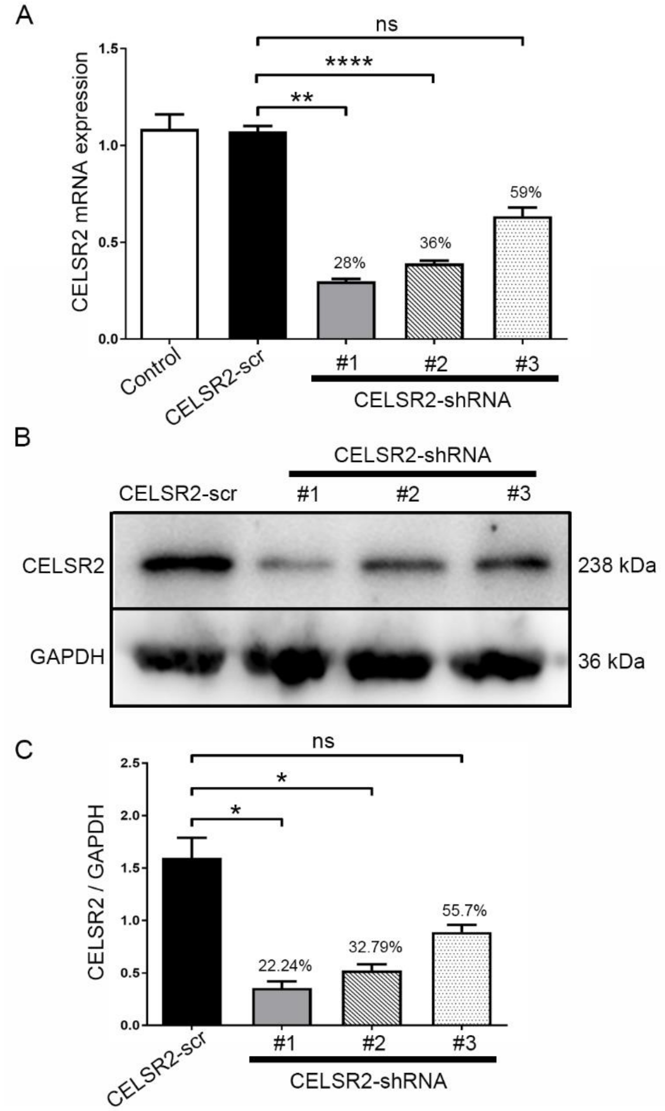


Cultured spinal motor neurons from WPC7 human embryos were transfected with CELSR2-scrambled shRNA (CELSR2-scr) or 3 different CELSR2-shRNA, and mRNA and proteins were extracted 3 days later.

(A) RT-qPCR showed CELSR2 expression in cultured spinal motor neurons with no transfection (Control) or after transfection of scrambled shRNA or CELSR2-shRNA. Using 3 different CELSR2-shRNAs, CELSR2 mRNA levels were 28%, 36% and 59% of control levels.

(B, C) Western blots with anti CELSR2 antibodies disclosed reduced protein levels in CELSR2-shRNA transfected cells (B). In cultured neurons transfected with 3 different

CELSR2-shRNAs (C), CELSR2 levels were 22.24%, 32.79% and 55.70% of those in control samples.

*, $P<0.05$; **, $P<0.01$; ****, $P<0.0001$; ns, not significant; Student $t$-test.

**Supplemental Figure 3. Human spinal motor neuron explants transfected by lentivirus.**

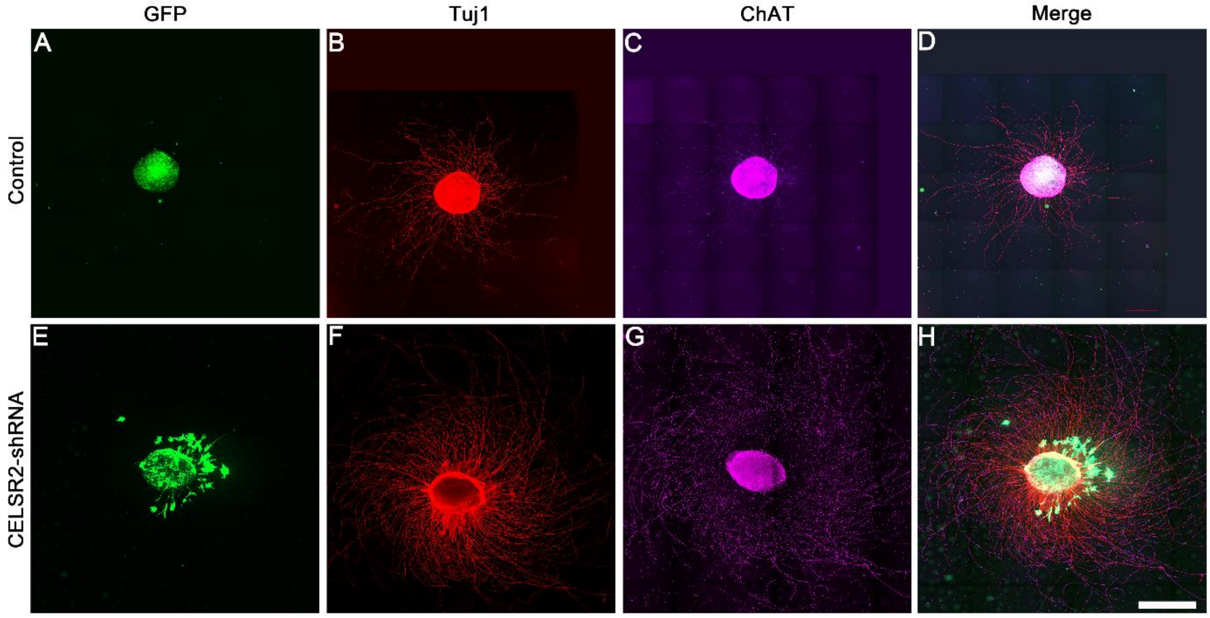


(A, E) Human spinal explants were transfected with scrambled shRNA-lentivirus (control, A) or CELSR2 shRNA-lentivirus (E). Lentivirus-encoded GFP labelled cells in the explants and some neurons that migrated out of the explants.

(B, C, F, G) After 5 DIV, cultured explants were immunostained with anti-Tuj1 (B, F) and anti-ChAT (C, G) antibodies.

(D, H) In the merged images, Tuj1- and -ChAT immunoreactivites overlapped in both groups, indicating that explants mainly include spinal motor neurons.

Scale bar: 500 µm.
